# Real-time decoding of bladder pressure from pelvic nerve activity

**DOI:** 10.1101/109942

**Authors:** Carl Lubba, Elie Mitrani, Jim Hokanson, Warren M. Grill, Simon R. Schultz

**Author notes:** These authors contributed equally to this study (Carl Lubba, Elie Mitrani).

## Abstract

Real time algorithms for decoding physiological signals from peripheral nerve recordings form an important component of closed loop bioelectronic medicine (electroceutical) systems. As a feasibility demonstration, we considered the problem of decoding bladder pressure from pelvic nerve electroneurograms. We extracted power spectral density of the nerve signal across a band optimised for Shannon Mutual Information, followed by linearization via piece-wise linear regression, and finally decoded signal reconstruction through optimal linear filtering. We demonstrate robust and effective reconstruction of bladder pressure, both prior to and following pharmacological manipulation.

## I. INTRODUCTION

There has been substantial recent interest in the development of bioelectronic medicines (also known as “electroceuticals”) [1]. Bioelectronic medicines consist of implantable devices capable of treating diseases by modulation of the nervous system. While substantial progress has been made in the treatment of some central nervous system disorders such as Parkinson’s Disease by deep brain stimulation [2], recently attention is focused on development of neuromodulation strategies for the peripheral nervous system [3]. The development of strategies for interfacing with and modulating the activity of peripheral nerves to the viscera may offer the prospect of extending bioelectronic medicine beyond diseases of the central nervous system, to the much larger class of non-neurological diseases that can be affected by electrical signalling in the peripheral nervous system, ranging from hypertension [4] to sleep apnea [5], rheumatoid arthritis [6] and sepsis [7].

Success will require advances in neural interfacing technology [8] and the development of closed loop peripheral neuromodulation devices. An important component of such a system is the ability to read out from the nerve, in real time, physiological signals of interest [9]. As well as providing an essential “dictionary” for modulation of peripheral nerves to control physiological variables, such readouts are an essential element of a closed loop control strategy to function in the presence of state changes and perturbations.

In the current work, we focus on the development of a decoder capable of reading out bladder pressure from electroneurograms recorded from the pelvic nerve of the rat. The pelvic nerve is a parasympathetic nerve that monitors and controls bladder function [10], and is thus an ideal testbed to evaluate PNS decoding algorithms. We show that by extracting a spectral signature optimised for mutual information conveyed about the pressure signal, linearizing to remove the effect of a static nonlinearity, and applying an optimal linear filter, it is possible to decode robustly and effectively bladder pressure from the pelvic nerve in both the Wistar and the Spontaneously Hypertensive Rat (SHR) model. We examined the effect on our decoder of delivery of PGE2, which generates acute symptoms of overactive bladder, and found that PGE2 resulted in systematic underestimation of bladder pressure by the decoder, which can be corrected by incorporating PGE2 data into the training set.

## II. METHODS

### A. Experiments

All animal care and procedures were reviewed and approved by the Institutional Animal Care and Use Committee at Duke University. Experimental data collection was similar to that described in detail in [11]. Briefly, adult Wistar (n=3) or Spontaneously Hypertensive Rats (n=3) were initially anaesthetized with isoflurane gas, followed by two subcutaneous injections of urethane totalling 1.2 g kg ^-1^. One hour after injections the animal’s reflexes were tested via foot pinch, and supplemental 0.1 g kg ^-1^ doses of urethane were given as needed until the foot withdrawal reflex abated. A cystometrogram comprising two to six bladder contractions was obtained by inserting a catheter into the bladder, and using a syringe pump connected to the catheter in series with a pressure transducer to pass room temperature 0.9% NaCl into the bladder at controlled flow rates. Recordings were made from the pelvic and hypogastric nerves, using nerve hook electrodes in mineral oil with 0.1% collagenase, and recorded with an ADInstruments Powerlab 8/35 system. Nerve signals were sampled at 10 kHz, and bladder pressure at 1 kHz.

### B. Frequency Band Selection

We extracted continuous signals in 50 Hz frequency bins from 1 Hz to 1 kHz without overlap, by using a Short Time Fourier Transform, with a window length of 0.5 s. The resulting signals were smoothed with a moving average window length of 10 s (for this analysis only), which was optimised to maximise mutual information while preserving temporal fidelity. To establish quantitatively the most informative frequency bands, we used the Kraskov estimator for the Shannon Mutual Information (MI) [12, 13] between the power spectral density for each band, and the bladder pressure time course. We selected the frequency range with MI above the mean for subsequent analysis, using the same range for all recordings.

**Fig. 1.**
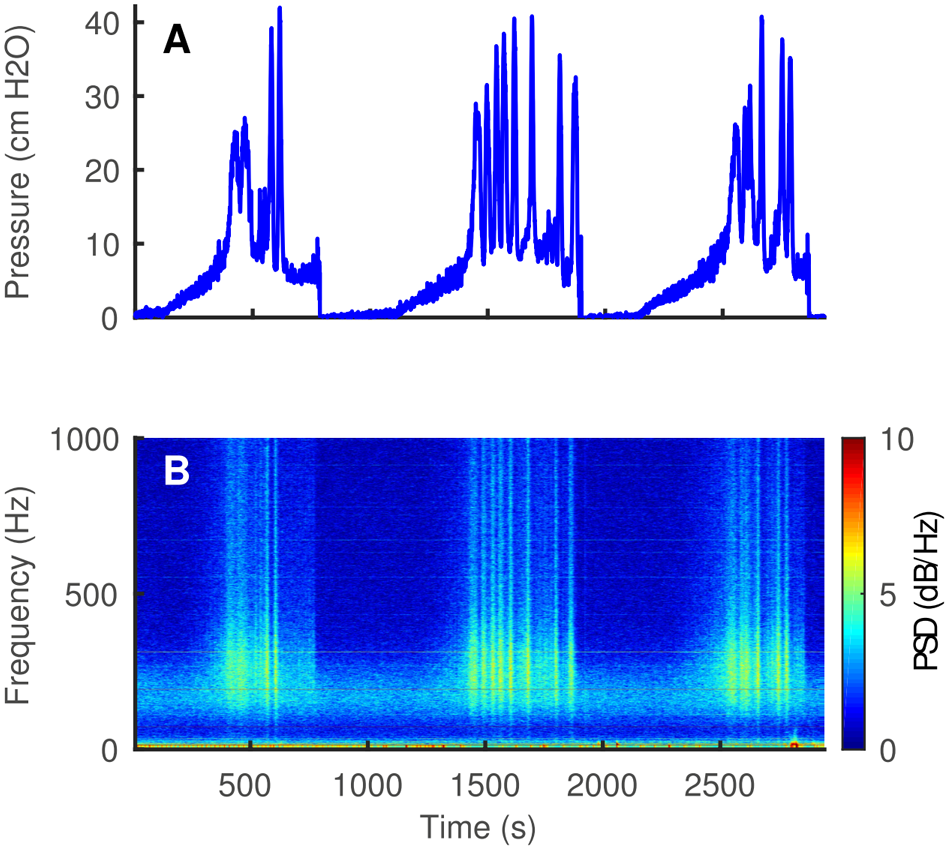
Cystometrogram. (A) An example pressure recording from a Wistar rat, showing three bladder contraction cycles. (B) Power spectral density (PSD) fluctuations throughout the contraction cycle.

**Fig. 2.**
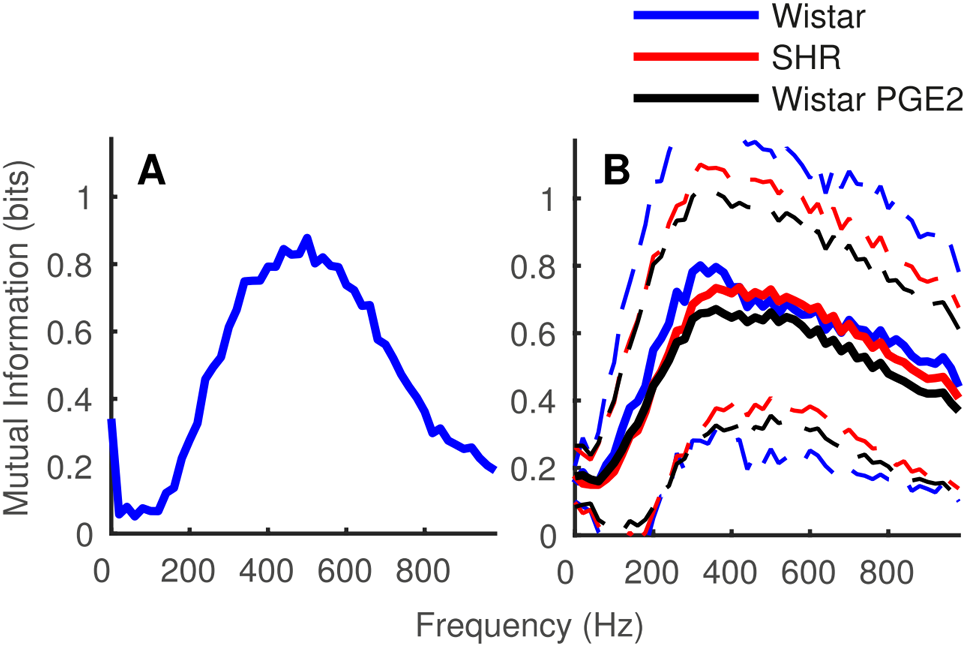
Mutual information between pelvic nerve activity and bladder pressure. (A) Information spectrogram for the example shown in Fig. 1. (B) Information spectrogram for the dataset of 3 Wistar and 3 SHR rats, as well as post delivery of PGE2 for two of the Wistar rats. Solid line: mean mutual information for the relevant frequency band; dashed lines indicate standard deviation.

### C. Piecewise Linear Fit

The simplest decoder we examined involved performing a piece-wise linear (PWL) fit of the instantaneous relationship between the measured pressure and the nerve power integrated across the spectral band from 200 Hz to 1 kHz (selected from the MI analysis). This PWL function was then used to look up the estimated pressure, instant by instant, for the test data.

### D. Optimal Linear Filter

An optimal linear filter (OLF) can build the time-dependent linear relationship between a series of measurements *x*_i_ and a quantity of interest *y*_i_ by convolving the last n measurement samples with a filter *β* and adding a constant offset. See 1. We use an OLF to derive the bladder pressure from the last *n* samples of nerve recording power in the selected frequency band.

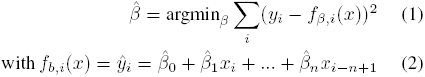

### E. Piecewise Linear Fit followed by Optimal Linear Filter

In this variant of the decoder, we used the training data to fit the nonlinear relationship between instantaneous nerve power in the 200 Hz to 1 kHz spectral band, and bladder pressure, and used that relationship to linearize the signal. The linearized signal was then decoded using an optimal linear filter, as described above, with filter coefficients regressed using training data.

### F. Cross-validation

Five-fold cross-validation was used throughout the analyses described here. Training data only was used for (i) the PWL fit, and (ii) recovery of optimal linear filter parameters. The MI analysis was instead used here to set generically the frequency band, identically, across all recordings. Performance was then measured on test data. For the training of the PWL model only, data was divided by an entirely randomized sample selection from the whole recording. For the OLF and combined models, the dataset was divided equally into sequential periods.

## III. RESULTS

### A. Information about bladder pressure is contained in a broad spectrum of pelvic nerve activity

We examined the frequency content of the pelvic nerve signal throughout the bladder contraction cycle, (Fig. 1), finding little power at low (< 200 Hz) frequencies, a band at moderate (200-400 Hz) frequencies that was present at low bladder pressures, but increased in power with bladder pressure, and rapid fluctuations in high frequency (400 Hz-1 kHz) power near the peak of the contraction cycle. To establish the best signal for decoding, we examined the MI conveyed about bladder pressure, by a narrowband signal scanned in frequency (Fig. 2). For the example shown in Fig. 1, MI about bladder pressure peaked at approximately 500 Hz (Fig. 2A). Across a population of 6 nerves, the spectral characteristics of the MI were broadly consistent, although typically peaking at a slightly lower frequency (< 400 Hz). The information spectrum did not differ between Wistar or SHR rats, nor did the delivery of PGE2 affect information content (Fig. 2B).

**Fig. 3.**
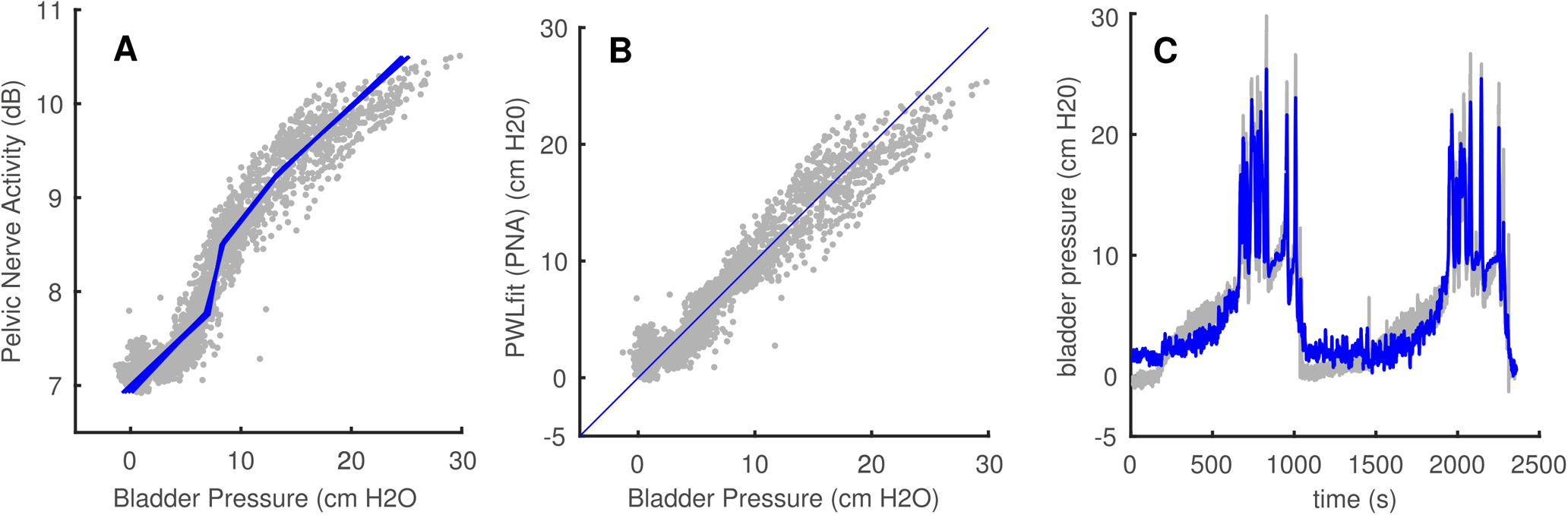
The relationship between bladder pressure and pelvic nerve activity can be fitted well as with a piecewise linear function (A). In our decoding approach, we use this fit to linearize the pressure-nerve activity characteristic (B). The result of the optimal linear filter on the linearized data over time follows the pressure reasonably well (C).

### B. Broad spectrum pelvic nerve activity provides a reliable but nonlinear representation of bladder pressure

A simple way to appreciate the relationship between bladder pressure and nerve activity is to plot the instantaneous time series values against each other (i.e. ignoring dynamics). This is shown in Fig. 4. Several things are immediately apparent: firstly, there is relatively little spread around the mean response to each bladder pressure value, indicating that the assumption of a lack of time-dependence is, at first order, reasonable. The residual noise may be accounted for by either noise or time dependence; some time-dependent effects may be addressed by the use of the linear filter approach below. A second observation is that the relationship follows a relatively stereotyped nonlinear form, differing however in detail for each nerve recorded (with respect to threshold, slope, etc). We take this to indicate that an instantaneous decoding approach should be relatively effective, with some small advantage potentially to be gained by the addition of a linear filter kernel.

**Fig. 4.**
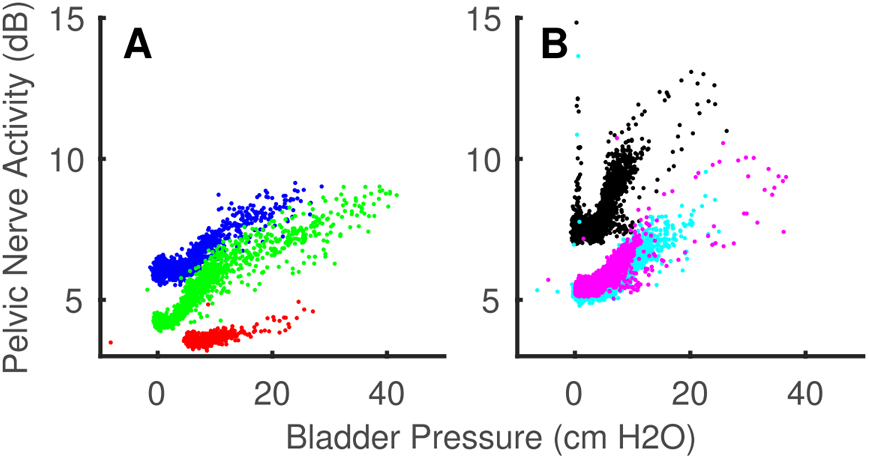
While nerve activity varies between individuals and type of rat (A: Wistar, B: SHR), all relationships between bladder pressure and pelvic nerve activity show a similar non-linear characteristic.

### C. Bladder pressure can be accurately decoded from pelvic nerve activity

We found that while reasonably good performance could be obtained by simply estimating the nonlinear relationship between nerve spectral power and bladder pressure, or by instead assuming linearity but incorporating time-dependence through an optimal linear filter, the best decoding performance was achieved by linearizing to remove the nonlinearity, then applying an optimal linear filter (see Table I). This approach assumes that there is a static nonlinearity, which is separable from the filter time-course. The PWL+OLF decoder was significantly better, in root mean squared error (RMSE) terms, than a PWL decoder alone (p=9.60e-4, one-sided Student’s t-test, n=6), and appeared to be slightly (but not significantly) improved over the OLF alone.

**TABLE I.** DECODER PERFORMANCE IN RMSE ± CROSS VALIDATION STANDARD DEVIATION (CM H2O)

| Rat | PWL | OLF | PWL + OLF |
| --- | --- | --- | --- |
| Wistar 1 | $2.26 \pm 0.10$ | $1.66 \pm 0.13$ | $1.62 \pm 0.19$ |
| Wistar 2 | $2.76 \pm 0.11$ | $2.31 \pm 0.40$ | $1.70 \pm 0.26$ |
| Wistar 3 | $2.49 \pm 0.23$ | $1.90 \pm 0.31$ | $1.96 \pm 0.29$ |
| Wistar (all) | $2.52 \pm 0.33$ | $1.96 \pm 0.28$ | $1.76 \pm 0.25$ |
| SHR 1 | $2.27 \pm 0.15$ | $2.06 \pm 0.45$ | $2.01 \pm 0.32$ |
| SHR 2 | $2.11 \pm 0.24$ | $1.65 \pm 0.18$ | $1.61 \pm 0.16$ |
| SHR 3 | $1.96 \pm 0.14$ | $1.27 \pm 0.48$ | $1.35 \pm 0.54$ |
| SHR (all) | $2.12 \pm 0.21$ | $1.66 \pm 0.37$ | $1.66 \pm 0.34$ |
| Wistar and SHR (all) | $2.32 \pm 0.20$ | $1.81 \pm 0.32$ | $1.71 \pm 0.29$ |

In the current version of the decoder, we used a simple PWL function to approximate the nonlinearity (Fig. 3A). The residuals from this model fit (Fig. 3B) suggest that the model can be improved, particularly at low bladder pressures. This leads to the decoder faithfully tracking the bladder pressure near the peak of the contraction cycle, but some systematic errors in the earlier phase of the contraction cycle (Fig. 3C), which should be addressable by incorporation of an improved, smooth, low order model function. The optimal linear filter function recovered (Fig. 5) includes structure extending 5-10 s back in time, which can be taken as an indication of the temporal fidelity of this system.

**Fig. 5.**
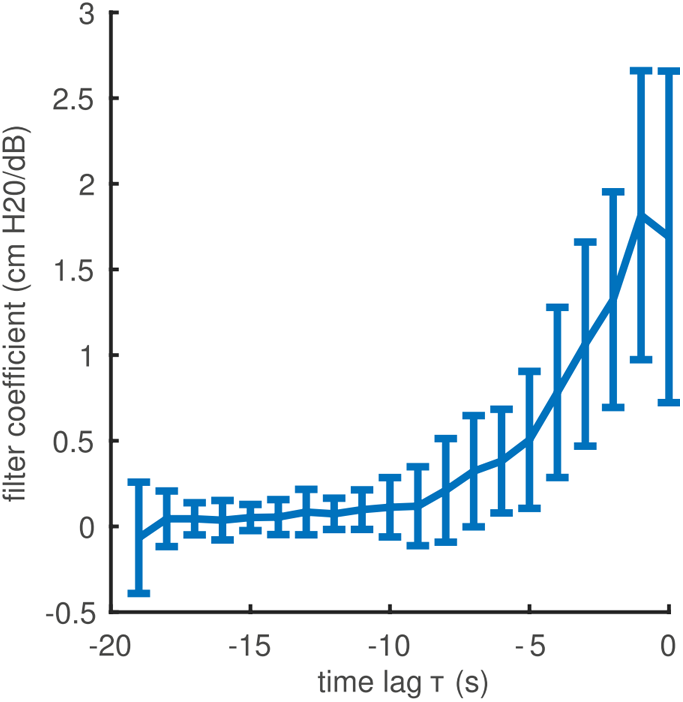
Filter coefficients recovered for the Optimal Linear Filter (mean values over n=6 Wistar and SHR rats, error bars indicate standard deviation).

### D. Decoding suggests that PGE2 administration leads to underestimation of bladder pressure by the pelvic nerve

Following recording of bladder pressure and nerve activity throughout the contraction cycle, in two experiments on Wistar rats, prostaglandin E2 (PGE2) was delivered into the bladder (100 µmol). This offered the opportunity to evaluate how the decoder would behave (i.e. what can be read out from the nerve) under a condition in which the system is pharmacologically perturbed from the conditions under which the decoder was trained. In both examples, the bladder pressure signal decoded from the nerve after PGE2 injection under-estimated the true bladder pressure (Fig. 6A). Training in the presence of PGE2 eliminates this bias (Fig. 6B).

**Fig. 6.**
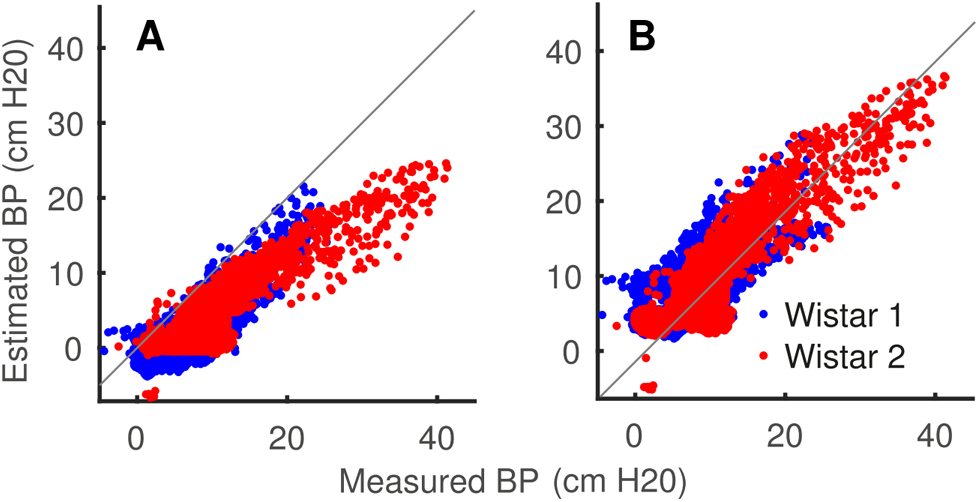
The administration of PGE2 causes the decoder to underestimate bladder pressure. (A) Mis-estimation due to training on pre- and testing on post-PGE2 administration data. (B) Performance achieved when training with PGE2 data (5-fold cross validation, same test data as A).

## ACKNOWLEDGMENT

We thank Daniel Chew, Kris Famm, Nick Jones, Victor Pikov, and Hannah Tipney for many useful discussions that led to this work.

